# Salicylic acid represses primary root growth through the Glucose-Target of Rapamycin-E2Fa pathway in Arabidopsis

**DOI:** 10.1101/2025.01.29.635412

**Authors:** Sanjay Singh Rawat, Ashverya Laxmi

## Abstract

In addition to their role as energy sources, sugars function as signaling molecules, modulating gene expression. Plants perceive nutrient availability and translate this information into cellular signals, to trigger various developmental responses, including primary root growth. In particular, glucose stimulates root development by activating the root meristem. Recent studies have documented the role of the defense hormone, salicylic acid (SA) as a negative regulator of root growth and developmental processes. Here, we characterized the modulation of primary root growth by the cross-talk of glucose and SA. Our results indicate that auxin is a critical mediator of SA-induced root growth inhibition. Attenuation of auxin signaling or transport pathways alters the inhibitory effect of SA on root growth. Moreover, we provide evidence for the involvement of the role of Target of Rapamycin (TOR) signaling in SA-induced root growth repression. Furthermore, SA negatively regulates the expression of E2Fa, a key transcription factor required for cell cycle progression and root growth and development. Our findings elucidate mechanism(s) whereby SA, through the interconnected glucose, auxin and TOR signaling pathways, inhibits primary root growth.

## Introduction

Glucose, in addition to its role as a primary energy source, functions as a versatile signaling molecule in plants (Sheen *et al*., 1999; Xiong and Sheen, 2013, 2015; Shi *et al*., 2018; Ryabova *et al*., 2019). It acts as a hormone-like substance, modulating various aspects of plant growth and development. Despite the existence of diverse sugar signals, glucose remains the most ancient and conserved, exerting control over various developmental programs by regulating gene and protein expression, cell cycle progression and metabolism (Xiong and Sheen, 2013, 2015; Rawat and Laxmi, 2024). Furthermore, glucose signaling has been shown to affect root growth and development, particularly primary root growth, lateral root (LR) development, LR orientation, gravitropism etc. by interacting with various phytohormones such as auxin, ethylene, gibberellins, cytokinin, brassinosteroids, jasmonic acid (JA) and abscisic acid (Zhou *et al*., 1998; Dekkers *et al*., 2008; Mishra *et al*., 2009; Kushwah and Laxmi, 2014, 2017; Gupta *et al*., 2015; Singh *et al*., 2015; Sharma *et al*., 2022). For instance, glucose alters the expression of diverse auxin signaling, biosynthetic as well as transporter genes and *glucose-insensitive* mutants such as the *gin2* are auxin resistant, establishing sugar and phytohormonal crosstalk in plants (Moore *et al*., 2003; Mishra *et al*., 2009, 2022; Sairanen *et al*., 2012).

The evolutionarily conserved serine/threonine kinase, Target of Rapamycin (TOR) acts as a pivotal regulator in plants, orchestrating cell proliferation and growth (Deprost *et al*., 2007; Dobrenel *et al*., 2016; Shi *et al*., 2018; Ryabova *et al*., 2019). By integrating diverse signals, including nutrients, energy and hormones, TOR enables plants to adapt to changing environmental conditions (Xiong and Sheen, 2015; Dobrenel *et al*., 2016). Specifically, TOR responds to sugars like glucose and the phytohormone auxin, synergistically promoting various aspects of plant growth and development (Xiong and Sheen, 2015; Dobrenel *et al*., 2016; Saxton and Sabatini, 2017; Schepetilnikov *et al*., 2017; Shi *et al*., 2018). The critical importance of sugars, in particular glucose, is evident in its necessity to activate TOR protein kinase, a process observed through the phosphorylation of its key downstream targets: S6 KINASE (S6K1), E2 PROMOTER BINDING FACTORa (E2Fa) and RIBOSOMAL PROTEIN S6 (RPS6) among several others (Xiong *et al*., 2013; Li *et al*., 2017; Chen *et al*., 2018; Shi *et al*., 2018). Glucose-TOR signaling constitutes a primary sugar signaling pathway that significantly influences cell proliferation within the root meristem, driving root growth and development (Xiong *et al*., 2013; Xiong and Sheen, 2013; Li *et al*., 2017). Remarkably, glucose-TOR directly phosphorylates E2Fa and promotes its activity in transcriptional activation of S-phase cell cycle genes (Xiong *et al*., 2013; Xiong and Sheen, 2013; Li *et al*., 2017). These findings uncover a novel role for glucose-TOR-E2fa in directly regulating gene transcription associated with the cell cycle, expanding beyond its established function of promoting protein synthesis through translational mechanisms through RPS6 that drive cell cycle progression (Mahfouz *et al*., 2006; Chen *et al*., 2018; Enganti *et al*., 2018). Within the Arabidopsis E2F family, three canonical members (E2Fa, E2Fb, and E2Fc) exhibit a strict requirement for dimerization with either Dimerization Partner (DPa) or DPb to facilitate robust and selective target DNA binding (Vandepoele *et al*., 2002; Del Pozo *et al*., 2005; Magyar *et al*., 2012). While E2Fa and E2Fb function as key activators of S-phase entry and progression, E2Fc diverges from its counterparts by lacking a transcriptional activation domain. Notably, E2Fc overexpression impedes cell cycle progression, whereas diminished E2Fc activity promotes cellular proliferation (Magyar *et al*., 2005; del Pozo *et al*., 2006; Polyn *et al*., 2015).

The phytohormone SA, not only is a well-known signal molecule mediating plant immunity, but is also involved in plant growth regulation. In Arabidopsis, SA is perceived by two classes of receptors: NONEXPRESSER OF PATHOGENESIS RELATED GENE 1 (NPR1) and NPR1-LIKE PROTEIN 3 (NPR3)/NPR4, which activate signaling pathways to stimulate the expression of defense-related genes and immunity (Fu *et al*., 2012; Peng *et al*., 2021; Jia *et al*., 2023). Since, the NPR family proteins do not contain DNA binding domains, NPR1 and NPR3/NPR4 interact with the transcription factors TGACG SEQUENCE-SPECIFIC BINDING PROTEIN (TGA) TGA2/TGA5/TGA6 for downstream signal transduction (Fu *et al*., 2012; Ding and Ding, 2020; Peng *et al*., 2021; Jia *et al*., 2023). Endogenous NPR1 localizes to both the nucleus and the cytosol and enhanced nuclear localization is critical to activate the transcription of *PATHOGENESIS-RELATED1*, (*PR1*) gene (Kinkema *et al*., 2000; Després *et al*., 2003; Pieterse and Van Loon, 2004; Peng *et al*., 2021). Recent research has consistently demonstrated that SA significantly influences root growth and development in a concentration-dependent manner. Low SA concentrations (<30 µM) have been shown to induce alterations in root architecture, such as the formation of the middle cortex and enlargement of the distal meristem. These effects are primarily driven by changes in auxin synthesis, transport and signaling (Wang *et al*., 2007; Pasternak *et al*., 2019). On the other hand, high exogenous SA induces root waving, regulates quiescent center (QC) cell division and stem cell maintenance and inhibits primary root growth, LR development as well as gravitropism (Zhao *et al*., 2015; Pasternak *et al*., 2019; Tan *et al*., 2020; Ke *et al*., 2021). In the present study, we describe the involvement of the auxin, glucose-TOR and E2Fa module in the repression of primary root growth by low levels of SA.

## Results

### 1) SA mediated regulation of primary root growth is sugar dependent

Previously, photosynthetic sugars like sucrose and glucose have been demonstrated to support primary root (PR) growth (Mishra *et al*., 2009; Xiong *et al*., 2013). To characterize the role of glucose-SA crosstalk in governing PR growth, 5-day-old seedlings of wild-type Columbia (Col-0) were grown onto ½ MS containing no sugar (control) and on glucose (166.52 mM) supplemented with increasing concentrations of SA. Plants grown on MS media supplemented with glucose displayed significantly longer PR lengths when compared to no sugar control (with or without SA) (**Figure 1a**). However, the progressive addition of SA at and concentrations exceeding 5 µM (15, 25, and 50 µM) resulted in a concomitant decrease in PR length in the presence of glucose, but not in its absence. This observation suggests that the inhibitory effect of SA on PR growth is contingent upon the availability of energy/glucose (**Figure 1a,b**). Furthermore, at relatively lower concentrations, SA did not affect the number as well as the length of the LRs, but induced a strong inhibitory effect on both these parameters at higher concentration i.e. 50 μM when compared to the control (**Figure 1c,d**).

**Figure 1.**
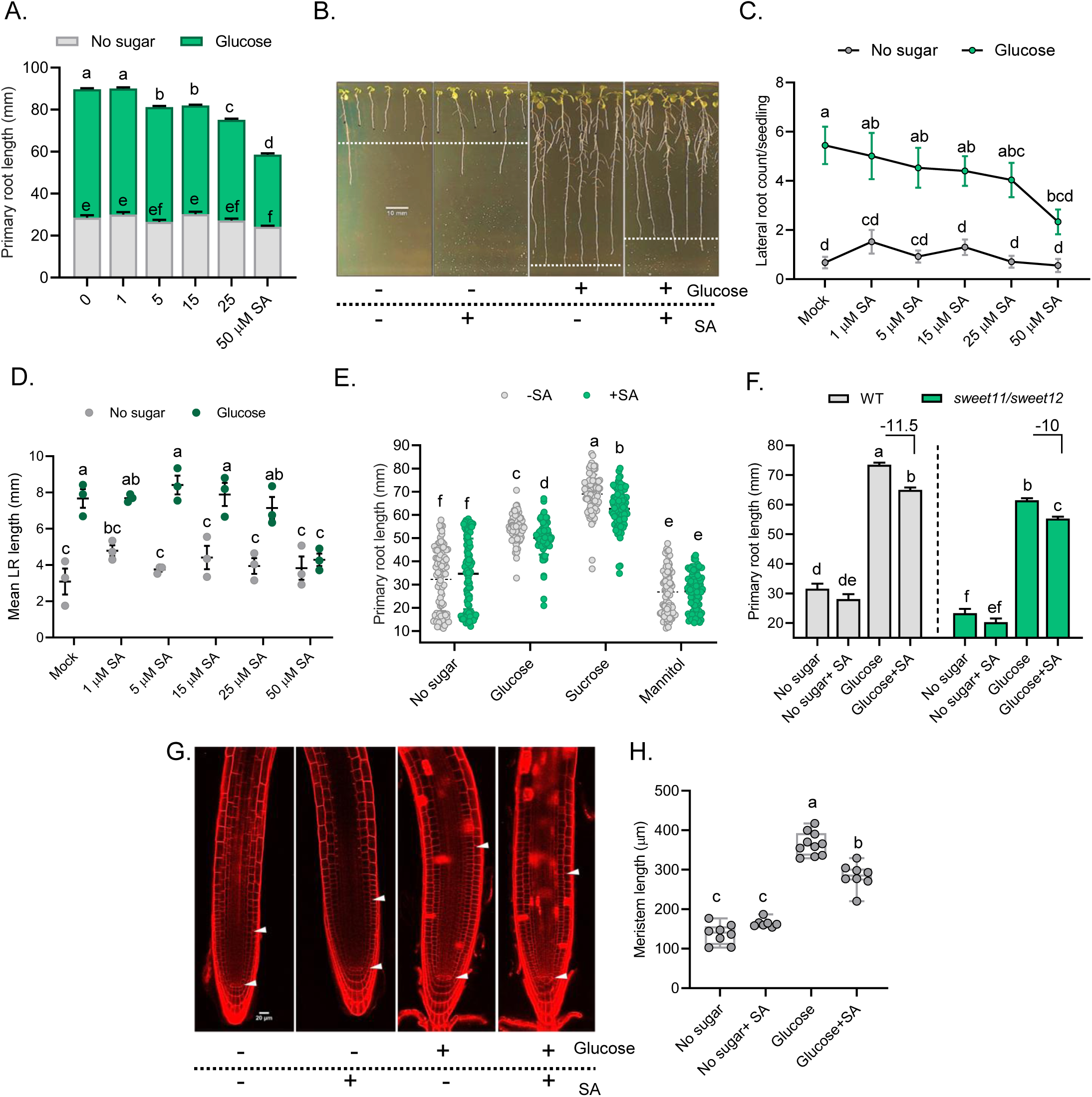
SA-mediated regulation of primary root growth is sugar dependent. **(A)** Analysis of PR growth of 12 day old seedlings after transfer to different doses of SA containing media, with and without glucose (n=90). Seedlings were photographed after 7 days from transfer. **(B)** Morphology of the seedlings pictured after 7 days from transfer on no sugar and glucose media supplemented with and without 5 µM SA. **(C)** Number of LRs of 12-d-old wild-type seedlings grown on ½ MS plate containing increasing concentrations of SA with and without glucose. **(D)** Mean LR length of seedlings grown on 1/2 MS plate containing increasing concentrations of SA with and without glucose (n=90). **(E)** PR lengths of seedlings grown on various sugars (n=90). **(F)** PR lengths of the WT Col-0 seedlings and the *sweet11/sweet12* mutant after treatment on different media (n=90). Numbers above the bars indicate percentage reduction in root length from glucose vs glucose plus SA treated roots. Experiments were repeated at least thrice yielding similar results. **(G)** Longitudinal confocal images of PI treated seedlings of Arabidopsis root meristems treated with SA grown on media with and without glucose for 48 hours (n=8). **(H)** Quantification of the meristem lengths of the seedlings in **(G)**. Experiments were repeated at least twice yielding similar results. Different letters indicate statistically significant differences between the means (one-way ANOVA followed by a post hoc Tukey’s honestly significant difference (HSD), *P < 0.05*).

To further elucidate the mechanism underlying the observed phenotype, 5-day-old wild-type seedlings were transferred to media supplemented with either metabolizable sugars (glucose and sucrose) or a non-metabolizable sugar (mannitol). We observed that SA retained its inhibitory effect on PR growth in the presence of both glucose and sucrose (**Figure 1e**). However, this effect was not observed in the presence of mannitol. These findings suggest that the inhibitory effect of SA on root growth, in conjunction with sugars, is not attributable to the osmotic effects of sugars, but rather to their metabolic effect (**Figure 1e**). Since sugar transport is a major determinant of overall plant growth, the root growth phenotype of the *sweet11, sweet12* mutant was analyzed under SA treatment. SWEET (<u>S</u>UGARS <u>W</u>ILL <u>E</u>VENTUALLY BE <u>E</u>XPORTED <u>T</u>RANSPORTER) transporters mediate the efflux of sugars like sucrose and hexoses like glucose and sucrose across various organs (Baker *et al*., 2012; Chen *et al*., 2015; Anjali *et al*., 2020). It was noticed that SA was able to inhibit PR growth to nearly same level as in the wild-type (**Figure 1f**), indicating sugar transport-independent role of SA on root growth inhibition.

Reportedly, glucose has been shown to activate and increase meristem size (Xiong *et al*., 2013) and given the established correlation between root meristem size and root length (Ubeda-Tomás *et al*., 2009; Hacham *et al*., 2011), we examined root meristems in wild-type Arabidopsis seedlings to determine if SA-mediated effects on root growth are constrained by meristem length. Therefore, we assessed the lengths of root meristems of seedlings grown under SA treatment with or without glucose. The root meristem length of SA-treated seedlings was found to be significantly shorter as compared to the roots treated with only glucose, suggesting that SA inhibits meristem growth to induce its effect on root growth (**Figure 1g,h**).

A closer examination of the root apical meristem was done next. The root stem cell niche (SCN) is composed of a group of infrequently dividing cells named the quiescent center (QC), surrounded by mitotically active stem cells, also called initials. Distal to the QC, the columella initials divide to form the mature differentiated columella cells (Dolan *et al*., 1993; van den Berg *et al*., 1995; Sarkar *et al*., 2007). Specifically, we observed that the overall number of columella cell layers (CSCs) increased from 4 to 5 (from no glucose vs. glucose-supplemented media) whereas the addition of SA to either glucose or to no sugar media did not result in any significant change (**Figure S1a,b**). WUSCHEL-RELATED HOMEOBOX 5 (WOX5) is a key transcription factor located in the QC, vital for stem cell maintenance and root development (Sarkar *et al*., 2007; Forzani *et al*., 2014; Kong *et al*., 2015). To assess the effect of glucose and SA on root meristematic activity, the *pWOX5::GFP* line was used (Sarkar *et al*., 2007). The intensity in the WOX5 expression was found to be maximum in the QCs of roots treated simultaneously with both glucose and SA (**Figure S1a,b**). The enhancement in WOX5 expression in the QCs corroborates with previous results (Wang *et al*., 2021). Previously, higher doses of SA have been shown to enhance the expression of WOX5 in extra tiers of the columella (Pasternak *et al*., 2019). However, the QC cell division nor the expression of WOX5 beyond the QC cells were observed under the SA concentrations used in this study as opposed to what has been reported by previous studies (Pasternak *et al*., 2019; Wang *et al*., 2021). This might have resulted from the usage of low concentration of SA. Overall, this suggests that low dose of SA leads to inhibition of root growth and this is modulated through energy dependent effect on the RAM.

### 2) AUX/IAAs mediate SA-repression of root growth

Previous studies have demonstrated that the biosynthesis of indole-3-acetic acid (IAA), the principal auxin in plants, is significantly upregulated by exogenous glucose and by physiologically relevant, low concentrations of the sugar (Mishra *et al*., 2009; Sairanen *et al*., 2012). These findings suggest that auxin signaling is a downstream process of glucose perception and signaling (Mishra *et al*., 2009; Sairanen *et al*., 2012). Several *auxin resistant* mutants such as *axr1*, *axr2*, *axr3* have been identified that exhibit pleiotropic growth phenotypes including their resistance towards auxin action (Lincoln *et al*., 1990; Leyser *et al*., 1996; Nagpal *et al*., 2000). These mutants have stabilized Aux/IAA proteins that are negative regulators of auxin signaling (Overvoorde *et al*., 2010). In line with the glucose and auxin crosstalk, findings have demonstrated that the auxin receptor *tir1* (*transport inhibitor response 1*) and response mutants *axr2*, *axr3* and *slr-1* (*solitary*-*root1*) not only exhibit defects in glucose-induced changes in root length, root hair elongation and LR production but also enhance the glucose-induced increase in root growth randomization from vertical (Mishra *et al*., 2009). This suggests that the effects of glucose on plant root growth and development are mediated by auxin signaling components (Mishra *et al*., 2009). We therefore analyzed whether SA application also leads to root growth inhibition in the auxin signaling mutants, since these mutants are perturbed in both glucose and auxin signaling. To this end, the PR lengths of two *auxin resistant* mutants namely; *axr1-3* and *axr2-1* were analyzed under glucose and SA treatments. We observed that the PR lengths of these mutants were more sensitive to SA than the wild-type seedlings (**Figure 2a**). In particular, the *axr1-3* and the *axr2-1* mutants exhibited 12.2% and 21.8% reduction in overall root growth, respectively compared to the wild-type (-8.7%) (**Figure 2a**). To further assess for the role of these repressors in SA-mediated root growth inhibition, the chemical PCIB (p-Chlorophenoxyisobutyric acid) which impairs auxin action in the roots via regulating the stabilization of Aux/IAA proteins was used (Oono *et al*., 2003). PCIB treated seedlings both at 1 and 5 µM, had shorter PR lengths than the no PCIB control upon SA treatment (**Figure 2b**). These results confirm SA negatively regulates root growth through the auxin signaling pathway.

**Figure 2.**
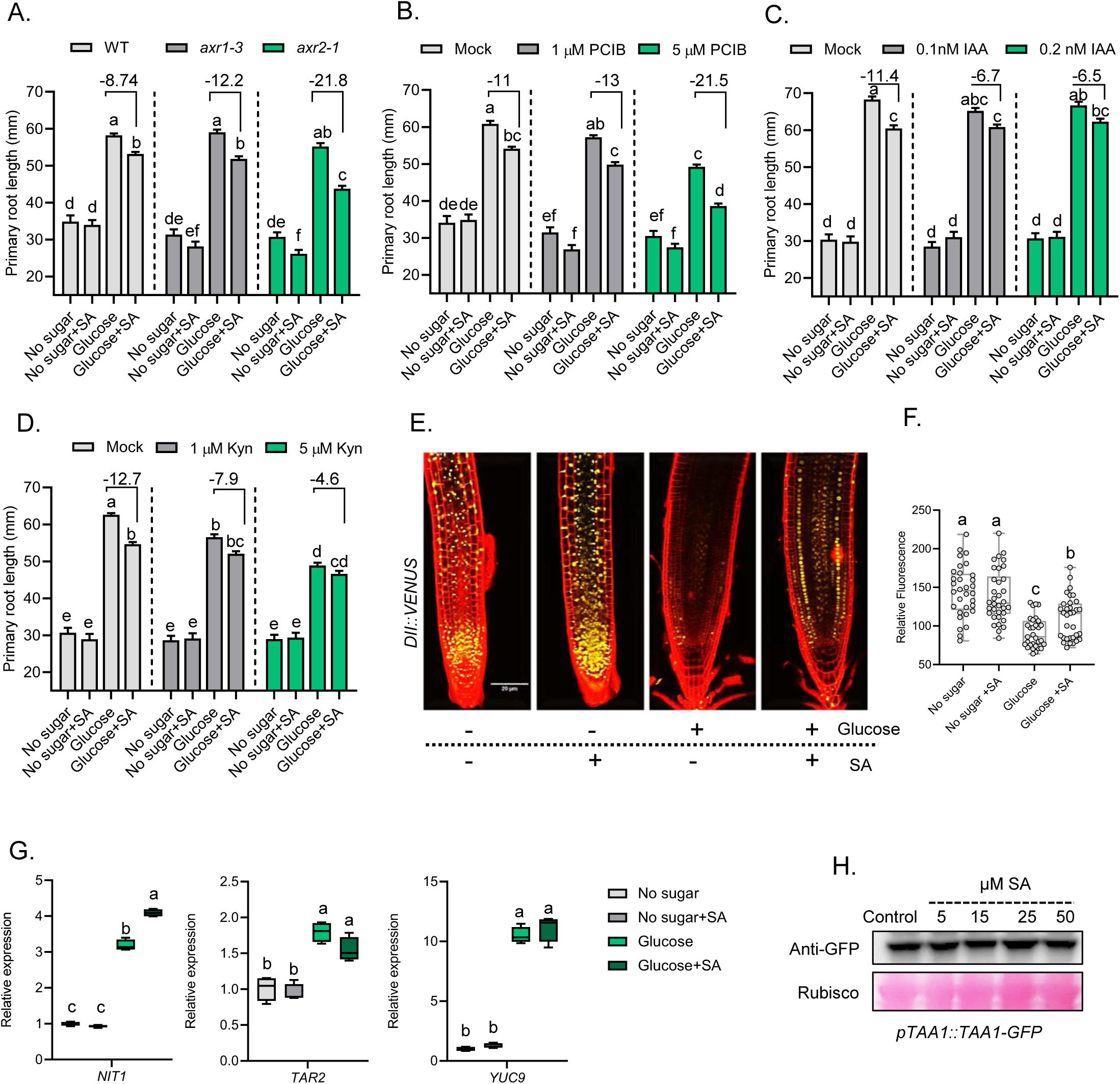
AUX/IAAs mediate SA-repression of root growth. **(A)** PR length of the auxin signaling mutants; *axr1-3* and *axr2-1* (n=90). **(B)** PR length of the WT Col-0 seedlings after treatment with auxin action inhibitor PCIB (n=90). **(C)** Effect of external auxin (IAA) application on WT Col-0 seedlings combined with glucose (-/+) and SA (-/+) treatments (n=90). **(D)** The effect of Kynurenine (Kyn) on WT Col-0 seedlings combined with glucose and SA treatments (n=90). **(E)** The effect of SA on the expression of *DII-VENUS* in the root tips of Arabidopsis seedlings. 5-day-old *DII-VENUS* expressing seedlings were grown on ½ strength MS after which they were transferred to MS media containing no sugar/glucose with or without SA for 48 hours. **(F)** Graphical representation of fluorescence intensities in figure **(E)**. **(G)** Relative expression of auxin biosynthetic genes under glucose and SA treatment. Experiments were repeated at least thrice yielding similar results. 5-day-old seedlings were starved for 24 hours before treatment with 5 µM SA with or without glucose for 6 hours. **(H)** Immunoblot assay of 5-day-old *pTAA1::GFP-TAA1* seedlings treated with varying concentrations of SA for 6 hours. Experiments were repeated at least twice yielding similar results. Different letters above the bars indicate statistically significant differences between the means (one-way ANOVA followed by a post hoc Tukey’s honestly significant difference (HSD), *P < 0.05*).

The above notion was further tested by performing root physiological experiments employing exogenous application of auxin (IAA). We found that the PR lengths of the IAA treated seedlings were slightly longer as compared to the no IAA control in the presence of SA at both the concentrations of IAA tested (0.1 and 0.2 nM), indicating that exogenous auxin can partially rescue the SA-mediated inhibition of root growth (**Figure 2c**). Conversely, the effect of the auxin biosynthesis inhibitor L-Kynurenine (Kyn) on root growth of wild-type seedlings was also tested. Kyn has been shown to inhibit PR growth in Arabidopsis seedlings (He *et al*., 2011). Biochemical and phenotypic analyses have revealed that Kyn competitively inhibits the activity of TRYPTOPHAN AMINOTRANSFERASE OF ARABIDOPSIS1/TRYPTOPHAN AMINOTRANSFERASE RELATEDs (TAA1/TAR), which are key enzymes involved in the indole-3-pyruvic acid pathway of auxin biosynthesis (Stepanova *et al*., 2008; He *et al*., 2011). In line with this, Kyn treatment mimics the loss of TAA1/TAR functions (He *et al*., 2011). We thus checked whether SA requires local auxin biosynthesis, which is important for root meristem activity (Stepanova *et al*., 2008; Brumos *et al*., 2018). At 1 µM Kyn, the ability of SA to repress PR growth was slightly repressed, however, at 5 µM Kyn, the PR lengths of SA treated seedlings were similar to their glucose grown control (**Figure 2d**). Based on these observations, we reasoned that SA-mediated root growth inhibition depends upon the presence of normal auxin gradient across the root meristem.

To explore whether auxin signaling also participates in SA-mediated inhibition of root development, we first assayed possible changes in auxin signaling in SA-treated roots using the auxin sensor *DII-VENUS* reporter which reflects upon internal auxin concentration (Brunoud *et al*., 2012). *DII-VENUS* is based on the DII domain of Aux/IAA28, expressed under a constitutive promoter and rapidly degraded in response to auxin (Brunoud *et al*., 2012). Previously, it was reported that high SA doses stabilize AXR2/IAA7 protein (Wang *et al*., 2007). In accordance, *DII-VENUS* activity was assessed in the presence of a high concentration of SA, specifically at 50 µM. Under high SA conditions, the fluorescence intensity of *DII-VENUS* expressing seedlings in the roots was elevated compared to the mock (**Figure S2a**). The *DII-VENUS* fluorescence intensities were also assessed under treatments of glucose and SA. Not surprisingly, the fluorescence intensity was lower in roots of seedlings grown on glucose media, presumably because glucose induces auxin biosynthesis (Sairanen *et al*., 2012) (**Figure 2e,f**). Moreover, our results show that roots of seedlings subjected to low SA concentration (5 µM), exhibited slightly increased *DII-VENUS* activity (**Figure 2e,f**). These data suggest that SA, even at lower concentrations tends to stabilize the activity of the auxin repressors, thereby inhibiting auxin signaling in roots.

Transcriptional expression analysis has indicated that SA leads to an increase in the transcript levels of the auxin biosynthetic gene *TAA1* (*TRYPTOPHAN AMINOTRANSFERASE OF ARABIDOPSIS1*) which encodes for a rate limiting enzyme involved in auxin biosynthesis from its precursor tryptophan (Stepanova *et al*., 2008; Brumos *et al*., 2018; Pasternak *et al*., 2019). We therefore investigated if comparatively lower dose of SA could also regulate auxin biosynthesis. The expression of genes that affect auxin levels, was examined first through quantitative real-time PCR in the presence of no sugar/glucose with or without SA supplementation. In particular, the expression of *NIT1* (NITRILASE1), *TAR2* (*TRYPTOPHAN AMINOTRANSFERASE RELATED2*) and *YUC9* (*YUCCA9*) was enhanced in the presence of glucose (**Figure 2g**). This was not surprising since, soluble sugars including glucose have been previously demonstrated to induce the expression of genes related to the production of endogenous auxin levels in Arabidopsis seedlings (Sairanen *et al*., 2012). However, except for *NIT1*, supplementation of SA did not result in further change in the expression of these genes, suggesting that SA probably does not interfere with auxin biosynthesis at the transcriptional level (**Figure 2g**). Furthermore, immunoblot assay involving SA treatment to Arabidopsis seedlings harboring *pTAA1::GFP-TAA1* transgene was done to analyze if SA was responsible for alteration in TAA1 activity. The result suggested that SA at all concentrations tested did not significantly lower the expression levels of TAA1 protein (**Figure 2h**). This suggests that at lower SA concentrations, the auxin biosynthetic pathway might be unaltered.

### 3) Auxin carriers mediate root growth inhibition upon SA supplementation

Polar auxin transport or PAT underlies auxin mediated root growth (Blilou *et al*., 2005). At the cellular level, various membrane localized transporters *viz.* PINFORMED (PIN), ATP-binding cassette group B (ABCB) efflux carriers and the uptake carriers AUX1/LAX (AUXIN1/LIKE-AUX) help mediate PAT (Kerr and Bennett, 2007; Swarup and Bennett, 2014). Apart from these, PID (PINOID) kinases and PP2A phosphatases maintain reversible PIN protein asymmetric distribution in cells to regulate root architectural and behavioral processes (Michniewicz *et al*., 2007; Armengot *et al*., 2014, 2016). To address SA requirement of a functional auxin transport, the PR lengths of the wild-type Col-0 seedlings were quantified after 1-N-naphthylphthalamic acid (NPA) and 2,3,5-Triiodobenzoic acid (TIBA) treatments, both inhibitors of auxin transport (Parry *et al*., 2001). We observed that at 0.5 µM NPA, the ability of SA to repress PR growth was unaffected, however, at 1µM NPA, the PR lengths of SA treated seedlings was similar to their glucose grown control (**Figure 3a**). Similarly, TIBA at a concentration of 2 µM was able to significantly repress the effect of SA on PR growth as compared to no TIBA control (**Figure 3b**). Additionally, to investigate whether the SA-mediated reduction of PR growth is dependent on auxin transport, the impact of mutations affecting auxin influx was examined. Compared to the wild-type (9.2%), both the *lax3* (4%) and *aux1-7* (6.6%) mutant showed less inhibition of PR growth when treated simultaneously with both glucose and SA (**Figure 3c**).

**Figure 3.**
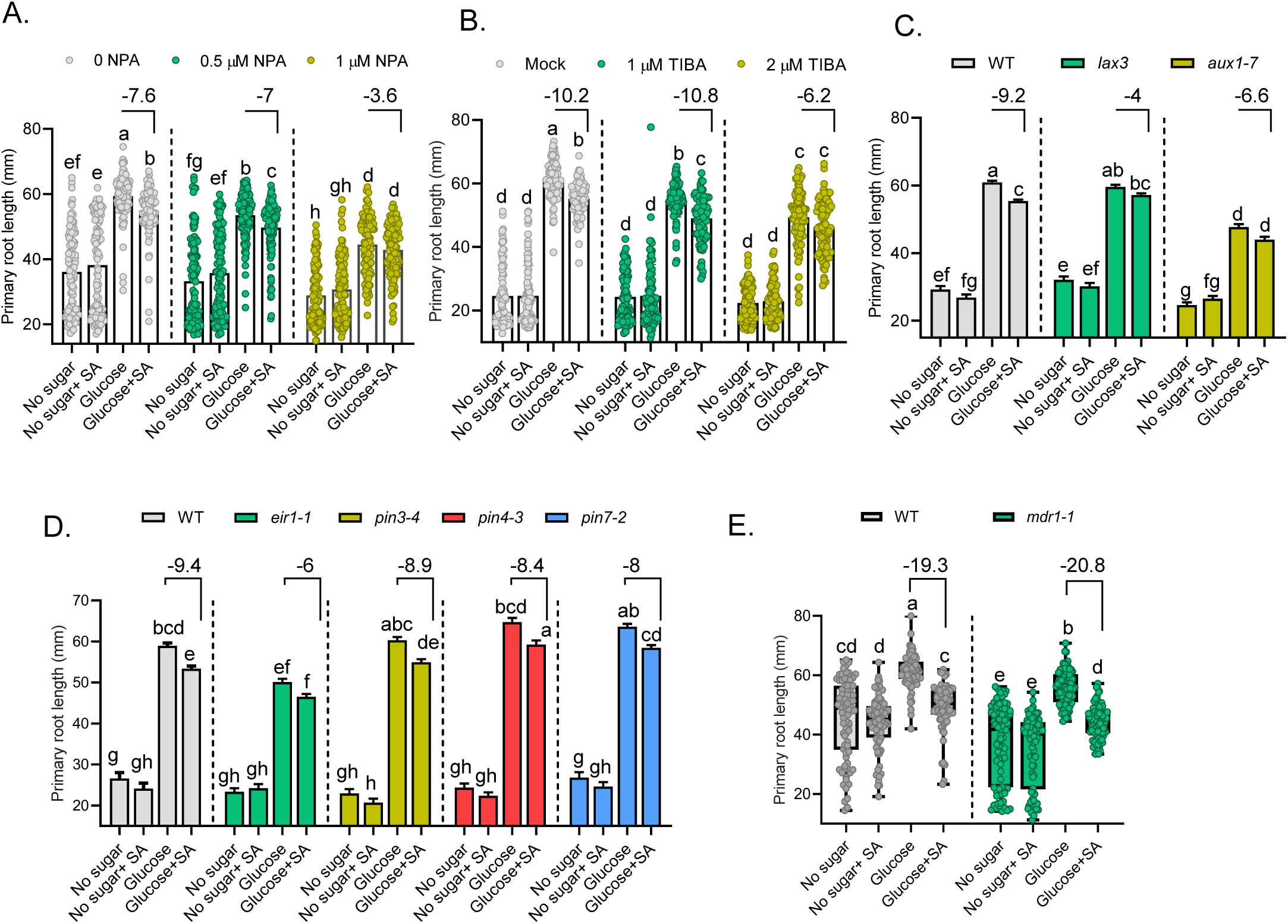
Auxin carriers mediate root growth inhibition upon SA supplementation. (A) The effect of the auxin transport inhibitor NPA on PR length grown on control/glucose-supplemented media with or without SA (n=114). (B) The effect of the auxin transport inhibitor TIBA on PR growth grown on control/glucose-supplemented media with or without SA (n=90). (C) PR length of the WT and the auxin influx mutants *lax3* and *aux1-7* grown on control/glucose-supplemented media with or without SA (n=115). (D) PR length of the auxin efflux mutants *pin2/eir1-1, pin3-4, pin4-3* and *pin7-2* grown on control/glucose-supplemented media with or without SA (n=95). (E) PR length of the auxin acropetal transport mutant; *mdr1-1* and its WT (Ws-2) (n=80). Seedlings were photographed on the 7^th^ day from transfer on indicated media. Experiments were repeated at least thrice yielding similar results. Different letters indicate statistically significant differences between the means (one-way ANOVA followed by a post hoc Tukey’s honestly significant difference (HSD), P < 0.05).

Next, the role of auxin efflux carriers in SA-mediated root growth inhibition was examined. Among these transporters, PINs play an important role in controlling root length (Friml *et al*., 2002; Feraru and Friml, 2008). In all the *pin* mutants studied, the roots of the (*ethylene insensitive root1*) *eir1-1*/*pin2* mutant exhibited the least reduction (∼6%) in terms of the SA-inhibition of PR length, while most other *pin* mutants *viz. pin3-4*, *pin4-3* and *pin7-2* exhibited somewhat similar response to SA as in the wild-type, suggesting that PIN3, PIN4, PIN7 might contribute redundantly to SA-mediated inhibition of root length, while PIN2 might have a predominant role (**Figure 3d**). Furthermore, the *mdr1-1* mutant allele, which is impaired in almost 80% root acropetal auxin transport (Lewis *et al*., 2007; Wu *et al*., 2007), exhibited wild-type-level sensitivity towards exogenous SA application on root growth (**Figure 3e**). In contrast, the roots of the *rcn1-1/pp2aa1* (*roots curl in naphthylphthalamic acid1*/ *protein phosphatase 2A)* mutant, which displays exaggerated basipetal auxin transport and defective PIN polarity (Rashotte *et al*., 2001; Kwak *et al*., 2002; Michniewicz *et al*., 2007; Sukumar *et al*., 2009), exhibited enhanced SA-mediated decrease in PR growth (**Figure S3a**). Previously, SA was shown to directly bind to the A subunit of PP2A, inhibiting activity of this complex (Tan *et al*., 2020). Since PP2A mediates the dephosphorylation of PIN2, PIN2 becomes hyperphosphorylated in response to SA (Tan *et al*., 2020). This hyperphosphorylation inhibits auxin transport and subsequently affects auxin-mediated root development, including growth, gravitropic response and lateral root organogenesis (Tan *et al*., 2020). Thus, the assays conducted on mutants such as *aux1*, *lax3*, *pin2*, and *rcn1*, which exhibit altered basipetal auxin transport, suggest that the inhibition of root growth by SA is primarily mediated by components of the basipetal auxin transport pathway.

The multifaceted interplay between ethylene and auxin intricately modulates the synthesis, signaling and transport of these hormones. Auxin upregulates *ACS* (1*-AMINOCYCLOPROPANE-1-CARBOXYLATE SYNTHASE*) transcription, thereby enhancing ethylene synthesis (Tsuchisaka and Theologis, 2004). Conversely, ethylene application stimulates the expression of auxin biosynthetic genes (Růžička *et al*., 2007; Stepanova *et al*., 2008). Moreover, root responses to ethylene are contingent upon basipetal auxin transport (Růžička *et al*., 2007) and the inhibition of auxin influx or efflux confers resistance to ethylene-induced root growth inhibition. We therefore tested the possibility of whether SA effect on root growth is also ethylene signaling dependent. To this end, the PR growth response of ethylene signaling and biosynthesis mutants was tested in the presence of SA. The *etr1-1* (*ethylene resistant1-1*), *ein2-1* (*ethylene insensitive2-1*) and *ein3-1* (*ethylene-insensitive3-1*) mutant exhibited similar root growth inhibition as in the wild-type (**Figure S3b**). In contrast, *eto1* (*ethylene overproduction1*), an ethylene-overproducing mutant which constitutively shows the ethylene-evoked triple reaction phenotypes in the absence of exogenously implemented hormone (Guzmán and Ecker, 1990), exhibited significantly reduced SA-mediated root growth inhibition (**Figure S3b**). This might be the ascribed to either increased ethylene-induced auxin biosynthesis or/and its transport. In particular, direct assessments of auxin levels have indicated its concentration increases at the root tip as a result of ethylene, hinting at ethylene’s role in stimulating auxin synthesis (Růžička *et al*., 2007). Furthermore, ethylene can also regulate the capacity of auxin transport (Negi *et al*., 2008). Based on these facts, it is speculated that the increased amount of auxin as well as its transport across the root in the ethylene overproducing mutant *eto1* might be sufficient to overcome the inhibitory effect of SA on root growth.

### 4) SA alters PIN activity across the root meristem

High SA levels have been previously reported to alter PIN expression dynamics to exert its effect on essential root growth processes and behavior such as root elongation, lateral root growth and gravitropism (Zhao *et al*., 2015; Pasternak *et al*., 2019; Tan *et al*., 2020). For instance, SA associates with a PP2A subunit, consequently inhibiting the enzyme’s phosphatase function on PIN proteins (Tan *et al*., 2020). This inhibition results in an accumulation of phosphorylated states of PIN2, a phenomenon known as hyperphosphorylation, which attenuates the proteins’ ability to facilitate auxin efflux, thereby impeding plant growth due to restricted auxin distribution (Ke *et al*., 2021). Furthermore, it has been demonstrated that external administration of SA and its inherent elevation in SA overproducing mutants both suppress the endocytic uptake of diverse proteins located in the plasma membrane, especially PIN1 and PIN2 (Du *et al*., 2013). In line with this, findings have also demonstrated that SA treatments at comparatively higher doses to Arabidopsis seedlings result in alteration in *PIN* transcriptional levels (Wang *et al*., 2007; Armengot *et al*., 2014; Pasternak *et al*., 2019). To this end, we first tested if SA at a lower concentration (5 µM) could also regulate PIN dynamics. Consequently, the impact of SA together with glucose on the auxin transporters at the transcriptional level was examined first through quantitative real-time PCR (**Figure 4a**). The expression levels of all the transporter genes except *LAX1* were found to be elevated in response to glucose treatment (**Figure 4a**). However, SA treatment resulted in the downregulation of these genes, especially *PIN1*, *PIN2*, *PIN3*, *PIN4* and *PIN7* (**Figure 4a**). These results are in accordance to what has been previously reported. Armengot and colleagues have demonstrated that exogenous SA can mediate transcriptional repression of most *PIN* genes (Armengot *et al*., 2014; Pasternak *et al*., 2019). However, besides PIN transporters, SA did not significantly affect the expression levels of *AUX1*, *LAX1* and *LAX2* (**Figure 4a**). These findings suggest that SA typically decreases the expression of PIN proteins, which in turn might adversely affect the regulation of auxin movement within the root system.

**Figure 4.**
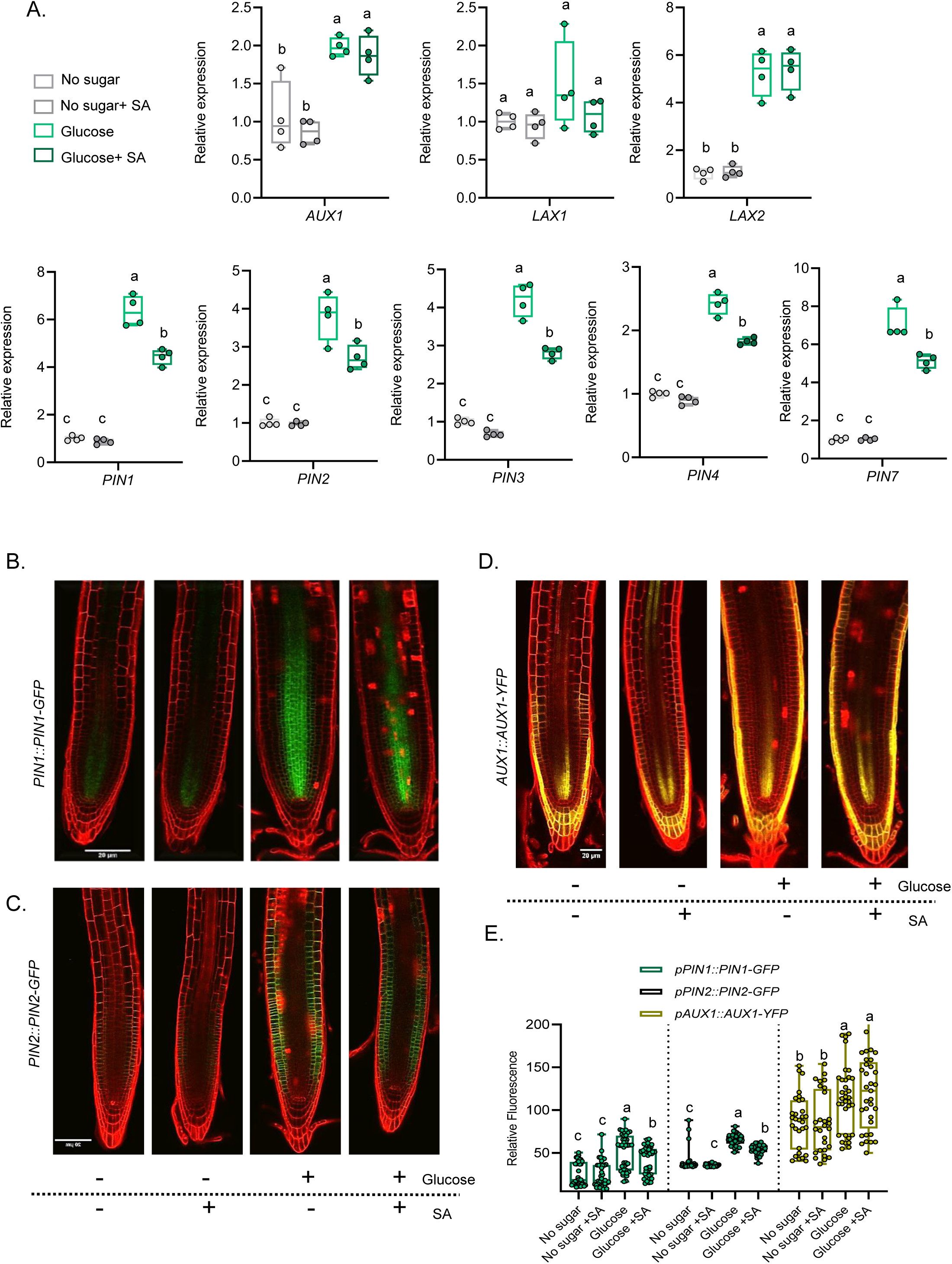
SA alters PIN activity across the root meristem. **(A)** Real-time qPCR analysis of the gene expression of auxin carriers in the WT seedlings treated with or without SA in combination with or without glucose. 5 day old seedlings were starved for 24 hours before treatment with either glucose and/or SA for 6 hours. Graphs are representative of one of three independent experiments yielding similar results (n=3). *UBQ10* was used as endogenous control. The expression levels of the indicated genes in the untreated roots (No sugar/control) were set to unity (1). **(B,C,D)** Expression of auxin efflux transporters in response to glucose and SA in primary roots. Arabidopsis transgenic *pPIN1::PIN1-GFP*, *pPIN2::PIN2-GFP* and *pAUX1::AUX1-YFP* seedlings were grown on control/glucose media supplemented with and without SA for 48 hours. Graphs in **(E)** illustrate differences in each reporter line expression, assessed as relative fluorescence intensity (n>28). Experiments were repeated at least thrice yielding similar results. Different letters indicate statistically significant differences between the means (one-way ANOVA followed by a post hoc Tukey’s honestly significant difference (HSD), P < 0.05).

Using GFP-tagged transgenic lines containing the promoters of PIN1, PIN2 and AUX1 i.e. *pPIN1::PIN1-GFP*, *pPIN2::PIN2-GFP* and *pAUX1::AUX1-YFP* protein fusions, we next examined whether SA affects PIN1, PIN2 and AUX1 protein localization. Our results indicate that the supplementation of glucose resulted in increased expression of all the transporters across the roots in comparison to the seedlings which were devoid of glucose (**Figure 4b,c,d,e**). Furthermore, SA treatment substantially reduced the levels of both PIN1 and PIN2 in the roots only when glucose was present in combination (**Figure 4b,c,d,e**). Moreover, despite the increased fluorescence intensity of the AUX1-YFP protein in seedlings grown in glucose-enriched media, the subsequent addition of SA did not significantly alter AUX1 distribution along the root meristem (**Figure 4b,c,d,e**). This result aligns with the findings previously reported by Tan *et al*., 2020. Together, these data suggest that SA-mediated root growth inhibition operates via interfering with PIN1 and PIN2 protein localization at the cell membrane, leading to the repression of auxin distribution across the root.

### 5) SA repression of root growth is TOR signaling dependent

Recent advances in glucose signal transduction research in Arabidopsis have revealed various pathways of glucose sensing: the HEXOKINASE1 (HXK1), *Arabidopsis thaliana* regulator of G-protein signaling (AtRGS1) and the TOR-mediated signaling pathways (Sheen *et al*., 1999; Moore *et al*., 2003; Urano *et al*., 2012; Shi *et al*., 2018; Ryabova *et al*., 2019). These pathways, through protein complex formation, orchestrate glucose signaling and metabolism, influencing diverse developmental processes (Jang *et al*., 1997; Sheen *et al*., 1999; Xiong and Sheen, 2013). To determine which sugar-signaling pathway mediates the sugar-dependent SA root growth inhibition, we tested the *hxk1-3* as well as the *tor RNAi* mutants, defective in glucose signal transduction (Moore *et al*., 2003; Deprost *et al*., 2007; Huang *et al*., 2015; Rawat *et al*., 2024). The *hxk1-3* mutant plants had only slightly but significantly shorter roots (-9.8%) on glucose plus SA medium compared to the wild-type (**Figure S4a**). In contrast, in the *tor RNAi*, the growth inhibition response to SA was greatly diminished (-1.2%) compared to wild-type (**Figure 5a**). Consistent with this, with 0.3 µM of AZD-8055, ATP-competitive TOR inhibitor (Montané and Menand, 2013), the PR length was significantly reduced relative to control plants (**Figure 5b**). On the contrary, no significant difference between AZD treated roots versus AZD plus SA treated roots was observed, suggesting that the presence of AZD eliminates the effect of SA on root growth inhibition (**Figure 5b**). These data indicate that TOR is essential for the SA-mediated PR root growth inhibition, while HXK1 has a very limited role in this. Since, SA leads to a reduction in the meristem size of the roots to exert its effect, we thus examined the effect of SA on the meristem size of both the mutant and the wild-type. In correlation with SA responsive reduction in meristem length of the wild-type, we determined that the meristem length of the mutant was not significantly affected by SA (**Figure 5c,d**).

**Figure 5.**
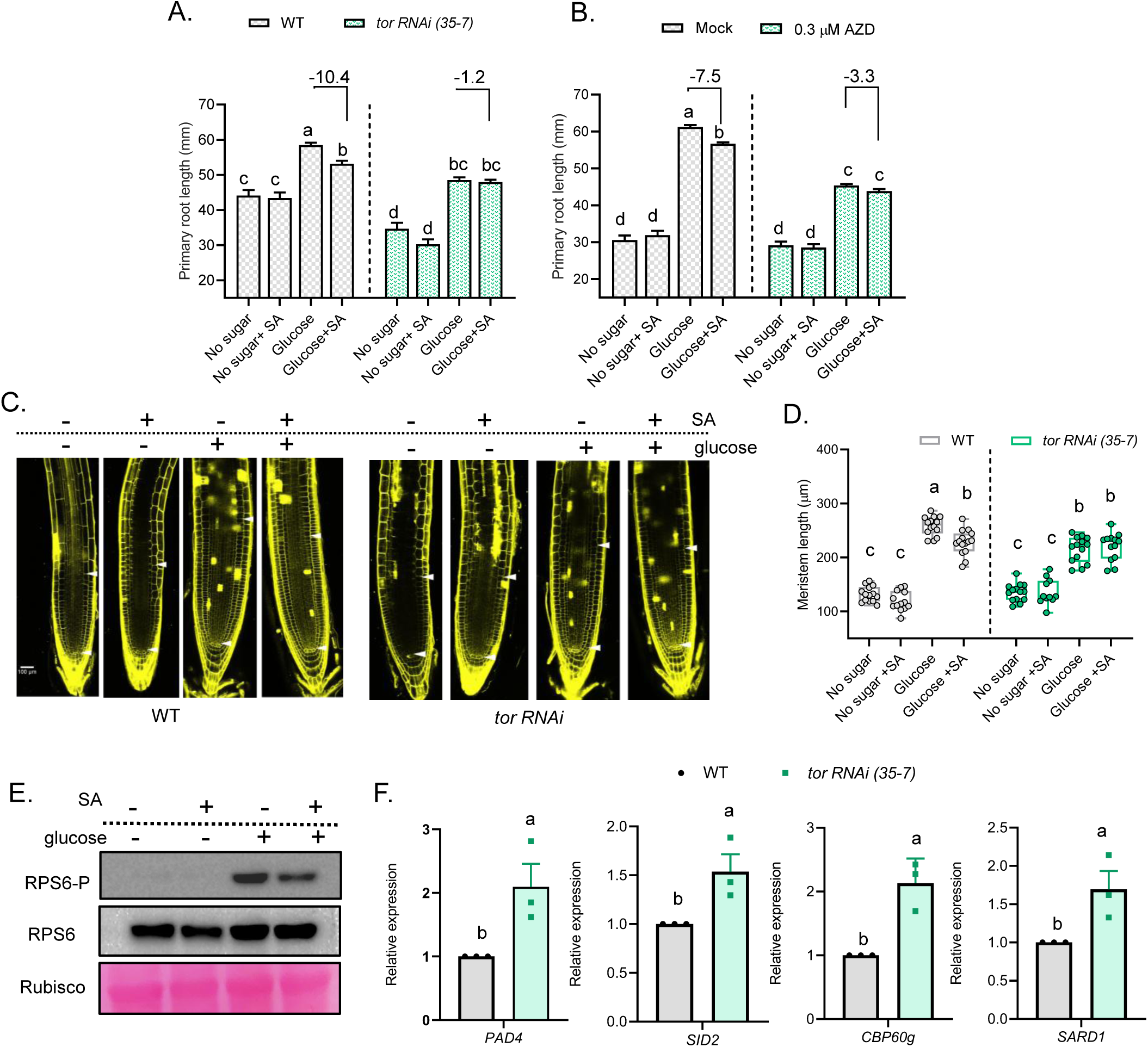
SA repression of root growth is TOR signaling dependent. **(A)** PR length of the WT and the *tor RNAi* mutant line (35-7) grown on no sugar/glucose media supplemented with or without SA (n=84). **(B)** The effect of the TOR inhibitor AZD-8055 on PR growth (n=112). Experiments were repeated at least thrice yielding similar results. **(C)** Longitudinal confocal images of WT and the *tor RNAi* mutant root meristems treated with SA grown on media with and without glucose for 48 hours. **(D)** Quantification of the meristem lengths of the seedlings (n>10) in **(C)**. Experiments were repeated at least twice yielding similar results (n=2). **(E)** Western blotting of the RPS6A-P (Ser240) and RPS6A proteins under glucose (-/+15mM glucose) and SA (-/+ 50µM SA) treatments. Seedlings were starved for 24 hours on no sugar MS media after which they were subjected to treatments indicated in the figure legends. Rubisco was taken as a loading control. Experiments were repeated at least twice yielding similar results. **(F)** Real time qPCR analysis of genes related to SA biosynthesis and signaling in WT versus the *tor RNAi* mutant line 35-7. *UBQ10* was used as an endogenous control. The expression levels of the indicated genes in the WT (control) were set to unity (1). Expression analyses were done thrice yielding similar results (n=3). Different letters indicate statistically significant differences between the means (one-way ANOVA followed by a post hoc Tukey’s honestly significant difference (HSD), P < 0.05).

Recent findings in various plants have indicated that pharmacological and genetic disruption of TOR strongly boosts SA and JA responses (De Vleesschauwer *et al*., 2018; Li *et al*., 2022; Marash *et al*., 2022). In particular, treating rice plants with rapamycin also resulted in a strong upregulation of the SA marker genes *NPR1* and *WRKY45*, suggesting that endogenous TOR antagonizes SA signaling (De Vleesschauwer *et al*., 2018). Hence, to gain more information about the effect of the SA on TOR signaling, we analyzed TOR activity *in vivo* through analyzing the phosphorylation of RPS6 (Dobrenel *et al*., 2016). Previous data has suggested that RPS6 is phosphorylated by the TOR kinase (Ruvinsky *et al*., 2005; Ren *et al*., 2012; Kim *et al*., 2014; Dobrenel *et al*., 2016). The degree of RPS6 phosphorylation also appears to be associated with increased protein synthesis and photosynthetic activities (Mahfouz *et al*., 2006; Turkina *et al*., 2011; Boex-Fontvieille *et al*., 2013; Dobrenel *et al*., 2016). We found that short term treatment of supplying exogenous glucose could effectively activate RPS6 phosphorylation (Ser240) compared with seedlings which were untreated/devoid of glucose (**Figure 5e**). Moreover, the application of SA led to a slight decrease in the phosphorylation of RPS6 (**Figure 5e**). This result suggests that SA might negatively affect TOR signaling components. Furthermore, to elaborate more on the TOR-SA antagonism, through expression analysis, we observed increased expression of various SA-signaling and biosynthetic genes including *PAD4* (*PHYTOALEXIN DEFICIENT4*), *SID2* (*SALICYLIC ACID-INDUCTION DEFICIENT2*), *CBP60g* (*CALMODULIN BINDING PROTEIN 60g*) and *SARD1* (*SAR DEFICIENT 1*) in the *tor RNAi* mutant seedlings when compared to the wild-type (**Figure 5f**). Together, the above data establish SA as a negative regulator of TOR-mediated signaling.

### 6) NPR1 signaling mediates SA-dependent PR growth inhibition

SA signaling in plant immunity is well known to be mediated by the NPR1, NPR3 and NPR4 receptors (Cao *et al*., 1997; Spoel *et al*., 2009; Peng *et al*., 2021). Hence, we examined the role of these receptors in SA-mediated root growth inhibition. We found, that the *npr1-2* mutant weakly exhibited the inhibition in PR growth by SA, while the decrease observed in *npr4-2* was similar to the wild-type (**Figure 6a**). These results are in contrast to previous reports where a NPR-independent mechanism of SA signaling has been shown to regulate various aspects of root growth and development (Tan *et al*., 2020; Ke *et al*., 2021). Furthermore, the higher order NPR mutant, *npr1*,*3*,*4* displayed a decrease in PR length similar to the wild-type when grown on SA, suggesting that the *npr1* mutation lies upstream of *npr3*,*4* mutations in regard to SA-mediated decrease in PR growth (**Figure S5a**). These results imply that SA-mediated PR growth inhibition in the presence of glucose is NPR1-dependent, while NPR3 and NPR4 play no significant role. To verify the effect of NPR1 on root growth, the constitutively NPR1 expressing *35S::NPR1-GFP* line, was studied next. When compared to the wild-type (7.6% decrease), the roots of the *35S::NPR1-GFP* seedlings displayed a significant response to SA treatment (10.5% decrease), suggesting that NPR1 acts as a positive regulator in mediating root growth inhibition by SA (**Figure 6b**). To further validate the hypothesis that exogenous SA-induced PR growth inhibition is a direct consequence of elevated endogenous SA levels, we utilized well-characterized mutants that constitutively overproduce SA, namely *snc1* (*suppressor of npr1*) and *cpr5-2* (*constitutive expresser of pathogenesis-related genes 5*) (Cao *et al*., 1994; Zhang *et al*., 2003). As expected, the PR lengths of these mutants were shorter compared to the wild-type when grown on only glucose supplemented media (**Figure 6c**). Moreover, SA supplementation together with glucose did not lead to any further decrease the PR length in these mutants, likely ascribing to already elevated endogenous SA levels (**Figure 6c**).

**Figure 6.**
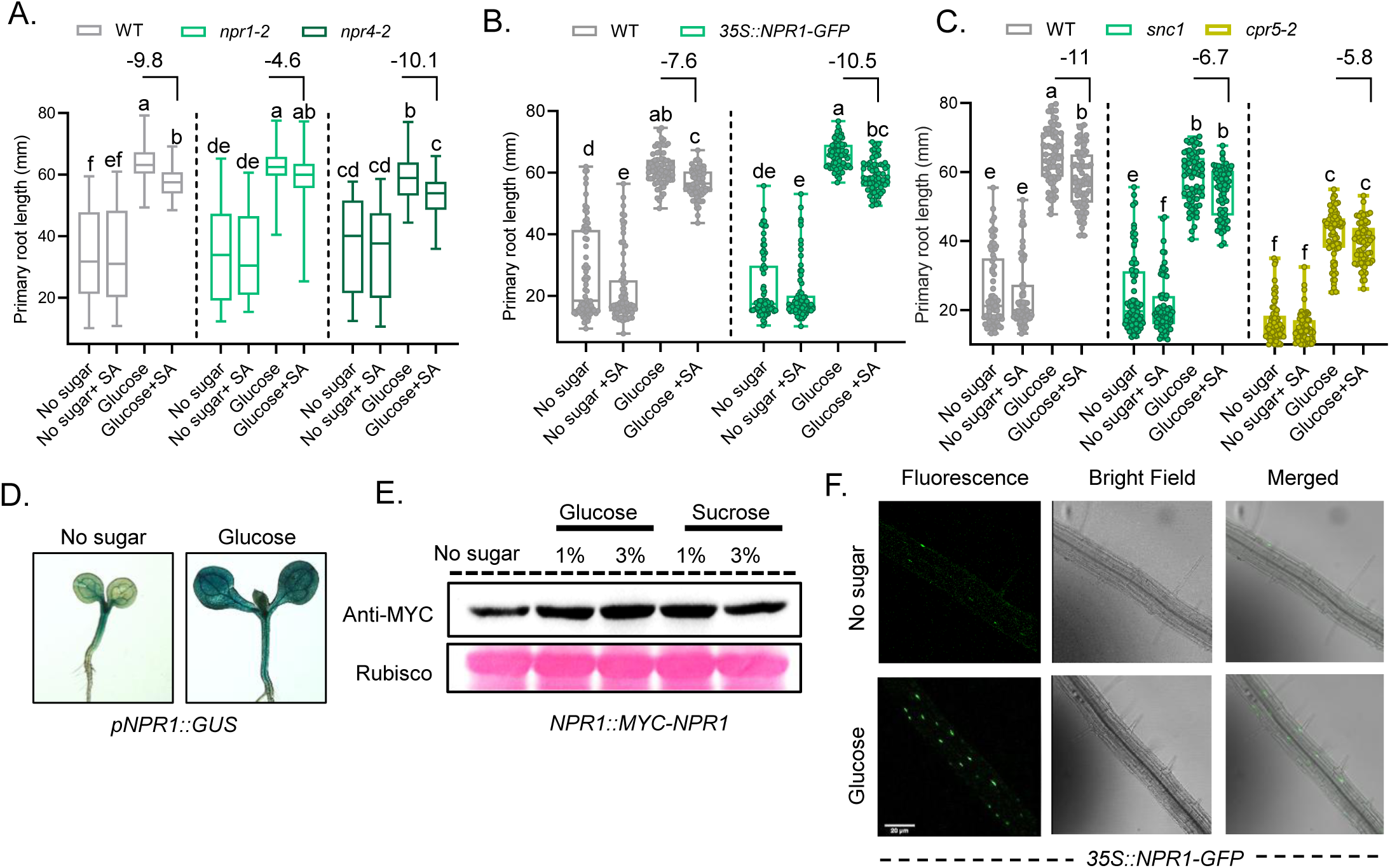
NPR1 signaling mediates SA-dependent PR growth inhibition. **(A)** PR length of 12-day-old seedlings with indicated genotypes of *npr* mutants (n=96). **(B)** PR lengths of WT Col-0 and *35S::NPR1-GFP* line grown on SA supplemented media with and without glucose. **(C)** PR lengths of SA overproducing mutants *snc1* and *cpr5* grown on glucose media with and without SA. **(D)** Seedlings expressing *NPR1::GUS* grown on control and glucose media for 48 hours. **(E)** 5-day-old *pNPR1::MYC-NPR1* transgenic plants were treated with varying concentrations of glucose and sucrose for 6 hours after 24 hours liquid starvation and subjected to western blotting using anti-Myc antibody. Rubisco was used as a loading control. **(F)** 5-day-old *35S::NPR1-GFP* expressing seedlings were grown on either no sugar or 3% glucose supplemented media for 48 h and were then imaged by confocal laser scanning microscope. Experiments were repeated twice yielding similar results. Different letters indicate statistically significant differences between the means (one-way ANOVA followed by a post hoc Tukey’s honestly significant difference (HSD), P < 0.05).

Various sugars, including sucrose, glucose and fructose, have been identified as signaling molecules in plant innate immunity. These sugars have been shown to upregulate the expression of *PATHOGENESIS-RELATED PROTEIN1* (*PR1*), a primary target of the NPR1 protein, in several plant species (Thibaud *et al*., 2004; Gómez-Ariza *et al*., 2007; Bolouri Moghaddam and Van den Ende, 2012). Interestingly, sorbitol, an osmotic agent, does not induce *PR1* expression, suggesting that osmotic effects of sugars are not involved in this process (Herbers *et al*., 1996). Recently, Yamada and Mine (2024) demonstrated that HXK1-mediated glucose signaling contributes to the enhancement of defense signaling during pathogen infection (Yamada and Mine, 2024). Therefore, in order to analyze the effect of glucose on SA mediated-NPR1 signaling, we first analyzed the expression pattern of the *pNPR1::GUS* line (Olate *et al*., 2018). Under glucose treated condition, a strong GUS activity was observed in the cotyledons of *pNPR1::GUS* line as opposed to untreated control (**Figure 6d**). This finding shows that the accumulation of *NPR1* mRNAs by glucose is regulated at the transcriptional level. We then tested the possibility if NPR1 protein levels are also regulated by glucose treatment. Consistent with the observed increase in *NPR1* transcript levels, immunoblotting analysis using the *NPR1::Myc-NPR1* line revealed that both 1% and 3% glucose as well as sucrose treatments significantly enhanced NPR1 protein levels when compared to the no sugar control (**Figure 6e**). Concomitant with this induction, we also observed that that the NPR1 localized more into the nucleus of the glucose-treated roots when compared to no sugar control (**Figure 6f**). This enhanced nuclear localization of NPR1 in response to glucose might lead to the suppression of growth-promoting genes (Li *et al*., 2016), a process that could be further amplified in conjunction with SA signaling. Collectively, our results suggest that glucose-induced increases in NPR1 levels prime plant roots for enhanced SA signaling, ultimately leading to reduced root growth.

### 7) SA treatment leads to inhibition of the expression of various genes related to S phase of the cell cycle

The TOR kinase, preserved throughout evolution in all eukaryotes, harmonizes signals from nutrients such as amino acids and energy, as well as growth factors in various organisms to orchestrate cell growth and the advancement of the cell cycle in a coordinated manner (Xiong *et al*., 2013; Shi *et al*., 2018; Rawat and Laxmi, 2024). Furthermore, the activation of TOR can stimulate the proliferation of root cells through the direct phosphorylation of E2Fa (Xiong *et al*., 2013; Li *et al*., 2017). This process subsequently boosts the expression of S-phase genes at the transcriptional level (Xiong *et al*., 2013; Xiong and Sheen, 2015). Firstly, in order to study the temporal expression of TOR, the accumulation of *AtTOR* transcripts was firstly analyzed by quantitative real-time PCR. Interestingly, no upregulation was observed in *TOR* expression upon glucose supplementation. Furthermore, the application of SA did not further alter the expression of *TOR* when compared to the control. This suggest that *TOR* might not be transcriptionally regulated by either short term glucose or SA treatments (**Figure 7a**).

**Figure 7.**
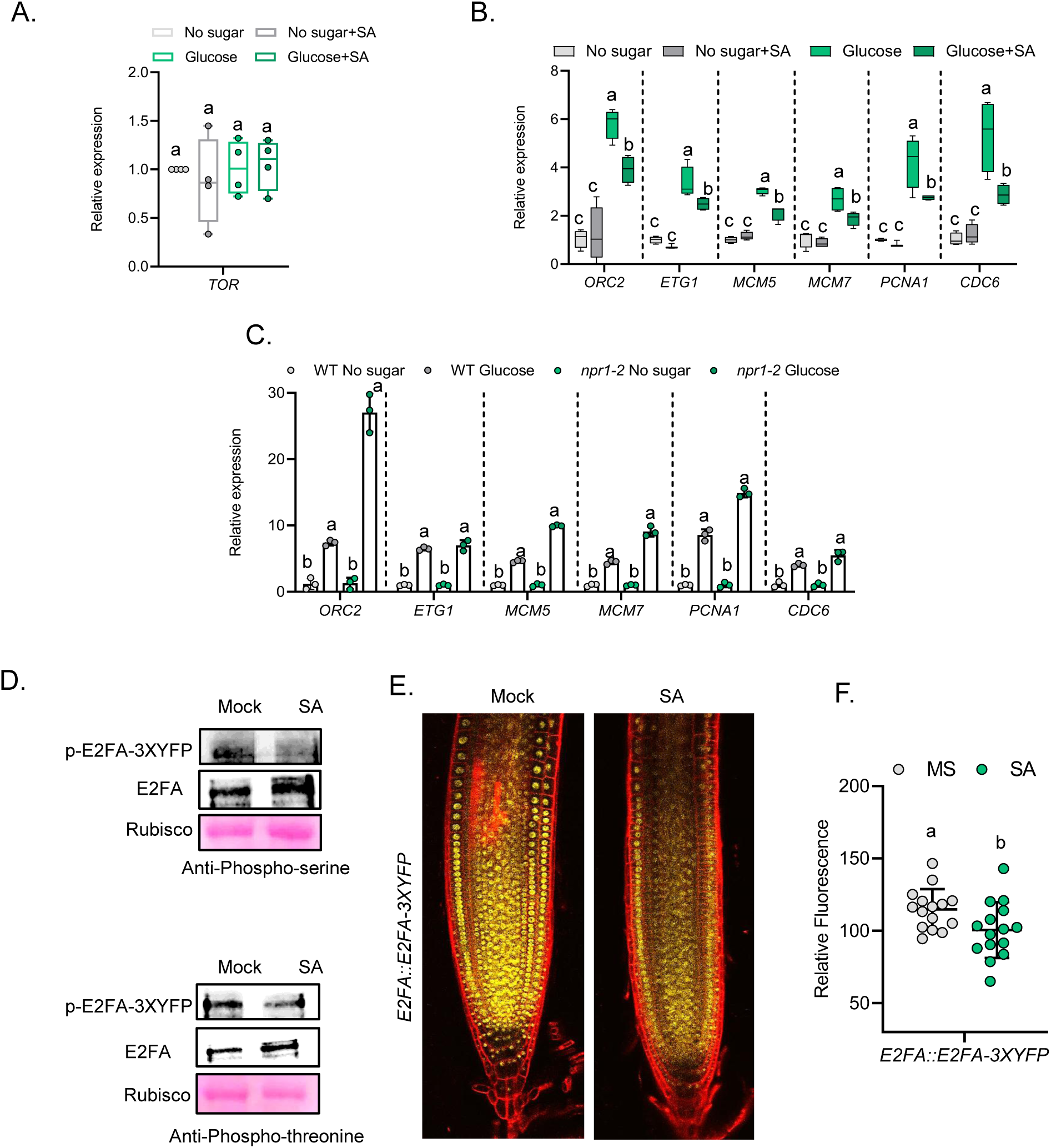
SA treatment leads to inhibition of the expression of various genes related to S phase of the cell cycle. **(A)** Real time qPCR analysis of *TOR* in WT upon various treatments. Experiments were done thrice yielding similar results (n=3) **(B)** GUS expression of the *CDC6a* in transgenic seedlings subjected to various treatments. *CDC6a::GUS* expressing seedlings were grown on control/glucose media with or without SA for 48 hours **(C)** Real time qPCR analysis of various S phase related genes in WT upon glucose/SA treatments (n=4). Seeds of the WT were sown directly on no glucose media after which they were grown for additional 5 days, followed by covering the media plates for 24 hours to eliminate any residual photosynthate. The seedlings were then subjected to treatments for 3 hours and the samples were harvested for expression analysis **(D)** Real time qPCR analysis of various S phase related genes in WT and *npr1-2* mutant (n=2). *UBQ10* was used as endogenous control. Experiments were repeated at least twice yielding similar results. **(E)** Western blot of immunoprecipitated E2Fa-E2Fa-3xYFP protein from seedlings grown on ½ MS media for 7 days with or without SA. Total proteins were extracted and cross-linked to the anti-GFP antibody and immunoprecipitation was performed using Protein-A/G coated beads. Some amount of rubisco was also immunoprecipitated along the target protein and was used as a loading control **(F)** Longitudinal confocal images of *E2FA::E2FAx3-YFP* expressing seedlings of Arabidopsis root meristems treated with and without 50 µM SA for 48 hours. **(G)** Quantification of the fluorescence intensities of the seedlings in **(F)**. Experiments were repeated at least twice yielding similar results. Different letters indicate statistically significant differences between the means (one-way ANOVA followed by a post hoc Tukey’s honestly significant difference (HSD), *P < 0.05*).

We next hypothesized whether SA might regulate cell cycle progression and for that the expression of S-phase-specific genes such as *CELL DIVISION CYCLE6* (*CDC6*), *ORIGIN RECOGNITION COMPLEX SECOND LARGEST SUBUNIT 2* (*ORC2*), *E2F TARGET GENE1* (*ETG1*), *PROLIFERATING CELLULAR NUCLEAR ANTIGEN1* (*PCNA1*), *MINICHROMOSOME MAINTENANCE3* (*MCM3*) and *MINICHROMOSOME MAINTENANCE5* (*MCM5*) (Takahashi *et al*., 2008; Xiong *et al*., 2013; Li *et al*., 2017) was examined in the wild-type Col-0 seedlings (**Figure 7c**). The transcription factors E2Fa/b are critical regulators that bind to promoter of these genes and determine the transition from the G1 to the S phase of the cell cycle (De Veylder *et al*., 2002, 2007; Friml *et al*., 2004; Li *et al*., 2017). Quantitative real time expression analysis indicated that the transcript level of these genes was elevated upon glucose supplementation but significantly decreased in the presence of SA (**Figure 7b**). Previous studies have indicated the promoter activity of the major cell cycle regulator, CDC6a is active in rapidly active meristems like root apices (Castellano *et al*., 2001), we asked whether SA treatment led to any change in the expression pattern of *CDC6a* in the root meristems. In contrast to those grown in the presence of glucose, the root meristems of the SA treated seedlings showed an attenuated expression of *CDC6a* activity (**Figure 7b**). However, no significant difference in the expression pattern of GUS in the absence of glucose with or without SA was observed (**Figure S6a**). We also checked whether the SA signaling mutant, *npr1* exhibited any alteration in the expression of these under glucose supplementation. Interestingly, the results revealed that E2Fa target genes were more upregulated in *npr1-2* compared to the wild-type, providing additional evidence for the functional interaction between glucose and SA signaling the regulation of cell cycle progression (**Figure 7c**). Furthermore, to investigate if the growth limitation of the primary root induced by SA signaling was a result of insufficient cell proliferation, the activity of cell division during the S-phase in the primary root meristems was observed using 5-Ethynyl-2′-deoxyuridine (EdU) staining. The EdU staining is frequently used as an indicator for the activity of cell cycle S-phase entry (Xiong *et al*., 2013; Li *et al*., 2017). Compared to the control (no sugar), the EdU staining signal was higher in the roots of the seedlings which were glucose treated (**Figure S6b**). Moreover, in accordance with the regulation of the cell cycle genes of the S phase by SA, the EdU fluorescence was also much higher in the *npr1-2* mutant than in the wild-type, suggesting that SA signaling negatively affects glucose-mediated enhancement in S-phase entry in the roots (**Figure S6b**).

Considering the essential role of the E2Fa in controlling cell division and differentiation during the G1/S cell cycle, we next examined whether E2Fa protein levels are also affected by SA. It has been suggested that phosphorylation events can increase E2Fs protein stability, which mediates enhanced binding on the promoter of S phase genes (Lin *et al*., 2001; Xiong *et al*., 2013; Li *et al*., 2017). To test this, we performed immunoprecipitation of the YFP labelled E2Fa protein with anti-GFP antibody on seedlings treated with or without 5 µM SA and then probed the immunoprecipitate with the both anti-phosphoserine and anti-phosphothreonine antibodies. The phosphorylated E2Fa protein levels were found to be significantly reduced in the presence of SA when compared to the control (**Figure 7d**). This indicates that SA application might lead to dephosphorylation of the E2Fa protein (**Figure 7d**). To investigate whether high SA doses could also result in the phosphorylation-dependent protein degradation of the pE2Fa::gE2Fa-3xYFP protein, E2FA fluorescence intensities were measured through confocal microscopy upon SA treatment (50µM) (**Figure 7e,f**). We found that the fluorescence signals in the roots were reduced significantly by SA. These results suggest that the E2Fa protein turnover is affected by both low as well as high doses of SA (**Figure 7e,f**).

To analyze if SA could also regulate the transition to G2/M phase, the *pCYCB1;1-CYCB1;1-GFP* reporter line was used to analyze the cell cycle progression in the root meristem. The expression of CYCB1;1 has been extensively used for monitoring cellular proliferation in various organs and is expressed in late G2 (Colón-Carmona *et al*., 1999; Schnittger and De Veylder, 2018). Enhanced cell divisions in the root meristem of glucose treated seedlings were observed as compared to the no sugar control (**Figure S6c,d,e**). This result aligns well with the previously established reports on glucose-activation of cell division in plant meristems (Skylar *et al*., 2011; Li *et al*., 2017). However, addition of SA with or without glucose, did not evoke any significant changes in the CYCB1;1 expression in the meristematic zone (**Figure S6c,d,e**). The above results also coincided with protein expression study through western blotting, suggesting that low SA levels might not regulate cell divisions to exert its effect on root growth and development (**Figure S6c,d,e**).

## Discussion

Plant growth, particularly primary root growth, is driven by interdependent actions of several growth-promoting hormones. Moreover, the crosstalk of these hormones with sugar signaling has been demonstrated to regulate a wide of root growth and developmental processes. The defense phytohormone SA, has emerged as a regulator of various crucial root growth processes (Pasternak *et al*., 2019; Kong *et al*., 2020; Tan *et al*., 2020; Ke *et al*., 2021; Wang *et al*., 2021). Here we show that the negative impact of SA on root growth repression is indeed energy dependent. Intriguingly, the inhibitory effect of SA on root growth is disrupted in the *npr1* mutant, unequivocally establishing NPR1 involvement in this process. Conversely, NPR3 and NPR4 appear to be dispensable for the SA response. We demonstrate that application of low levels of exogenous SA (5 µM) caused root meristem inhibition by dramatically reducing the root meristem size. Remarkably, this effect is independent of glucose transport, likely due to the extensive repertoire of sugar transporter genes in vascular plant genomes, encoding for a diverse array of sugar carriers (Reinders, 2012; Pommerrenig *et al*., 2020). Furthermore, we showed that auxin plays critical roles during SA inhibition of root growth. Previous studies have indicated that both auxin signaling as well as its biosynthesis are greatly affected by the presence of glucose (Mishra *et al*., 2009, 2022; Sairanen *et al*., 2012). Glucose and auxin exhibit coordinated regulation of root growth. For instance, the glucose-insensitive mutant *gin2* display reduced auxin sensitivity (Moore *et al*., 2003). Conversely, auxin-related mutants *axr1*, *axr2*, and *tir1* are glucose-insensitive (Mishra *et al*., 2009). Furthermore, genome-wide transcriptional profiling reveals glucose-mediated regulation of auxin-related genes, including biosynthetic enzymes, transporters, receptors and signaling molecules (Mishra *et al*., 2009). Therefore, we analyzed the role of glucose-mediated auxin signaling in the regulation of root growth inhibition by SA. In order to further evaluate the cross talk between auxin and SA in controlling root elongation, we examined the growth of mutants affected in auxin signaling as well as transport. Both pharmacological and mutant analyses indicate that SA is negatively involved in both auxin signaling and uptake. In particular, our results indicate that mutants affected in auxin signal transduction (*axr1-3*, *axr2-1*) show enhanced sensitivity towards SA, while those affected in auxin transport (*lax3*, *aux1-7*, *eir1*) exhibited reduced response. Moreover, the results of our *DII-VENUS* protein stability assay suggest that SA at both low as well as high doses, prevents the auxin-dependent degradation of the AUX/IAAs protein probably through the ubiquitin proteasome pathway (Wang *et al*., 2007). A similar working principle for SA to repress growth is linked through the repression of gibberellin pathway (Yu *et al*., 2022). The previous study revealed that NPR1 functions as an adaptor of ubiquitin E3 ligase to promote the polyubiquitination and degradation of GID1, which enhances the stability of DELLA proteins, the negative regulators of GA signaling (Yu *et al*., 2022). Given that hormones like cytokinin and brassinosteroids also stimulate root growth alongside auxin and gibberellin, it is crucial to examine whether SA also exerts its growth-inhibitory effects by interfering with other hormonal signaling pathways.

In addition to its role in auxin biosynthesis, glucose is also implicated in auxin distribution (Mishra *et al*., 2009) and our results show that SA application negatively regulates this process. In this study, we found that *PIN* genes are downregulation by SA. Moreover, we have demonstrated that glucose was able to induce the plasma membrane localization of the efflux transporters, namely PIN1, PIN2 and AUX1 and that SA was able to reduce their abundance, although no change in the AUX1 protein expression was observed. A reduction in the capacity of auxin transport to the root apical meristem could lead to a global decrease in the recirculation of auxin, explaining the reduction in root growth.

Finally, we show the glucose signaling is essential for glucose-induced root growth. To elucidate the sugar signaling pathways involved in SA-mediated growth repression, we employed HXK1-deficient mutants and *tor RNAi* plants. Our findings revealed that both pathways are essential for SA-induced inhibition of root growth, with the latter potentially playing a more prominent role. Furthermore, this impaired repression of SA on root growth was mimicked by exogenous treatment of with the TOR kinase inhibitor AZD8055. Interestingly, the crosstalk between TOR signaling and defense signaling has been documented in recent studies. Studies involving the TOR-specific inhibitor rapamycin, have revealed that TOR not only dictates transcriptional reprogramming of extensive gene involved in central and secondary metabolism, cell cycle and transcription but also suppresses many defense-related genes (Xiong *et al*., 2013; Li *et al*., 2017, 2022; De Vleesschauwer *et al*., 2018; Marash *et al*., 2022). On the one hand, TOR overexpression lines display increased susceptibility to both bacterial and fungal pathogens, whereas plants with reduced TOR signaling show enhanced resistance (De Vleesschauwer *et al*., 2018; Marash *et al*., 2022). Conversely, silencing of TOR has been shown to activate a subset of defense-related genes and promote resistance against several pathogens including *Pseudomonas syringae*, *Alternaria alternata*, *Xanthomonas euvesicatoria* and *Botrytis cinerea* (Meteignier *et al*., 2017; De Vleesschauwer *et al*., 2018; Marash *et al*., 2022). Interestingly, SA showed potent inhibitory activity against fungal pathogen*, Fusarium sp.* by targeting the TOR signaling pathway. In particular, SA was shown to activate the AMPK (AMP-ACTIVATED PROTEIN KINASE) ortholog SNF1 (SUCROSE NON-FERMENTING 1) in *Fusarium* to counter TOR signaling (Li *et al*., 2022). Earlier studies have shown that AMPK phosphorylates the TOR subunit RAPTOR to inhibit growth promoting activities (Gwinn *et al*., 2008; Nukarinen *et al*., 2016). We detected the phosphorylation of AtRPS6, which is the output of TOR signaling and is essential for protein synthesis and cell cycle progression, to be lowered upon SA treatment, further confirming that SA negatively influences TOR signaling. In line with this, we observed that SA-biosynthetic and signaling genes were also upregulated in the *tor RNAi* mutant. These data point towards the role of TOR-SA antagonism in Arabidopsis. Moreover, the glucose/sucrose activation of TOR kinase has been demonstrated to facilitate E2Fa-dependent transcriptional activation of S-phase genes in root meristem development (Xiong *et al*., 2013; Li *et al*., 2017). In particular, E2Fa phosphorylation is known to enhance its transcriptional activity. We show that SA-inhibition of root growth may also depend on an E2Fa-regulated cell cycle process. Indeed, E2Fa protein stability was seen to be decreased on both low as well high SA dose application. Consequently, quantitative real-time PCR analysis showed that transcript levels of S-phase related genes were downregulated upon exogenous SA treatment, which was in line with the upregulation of these genes in the SA signaling mutant, *npr1*. In conclusion, this study indicates that SA-dependent root growth inhibition is mainly regulated by the glucose-auxin-TOR-E2Fa module **(Figure 8)**.

**Figure 8.**
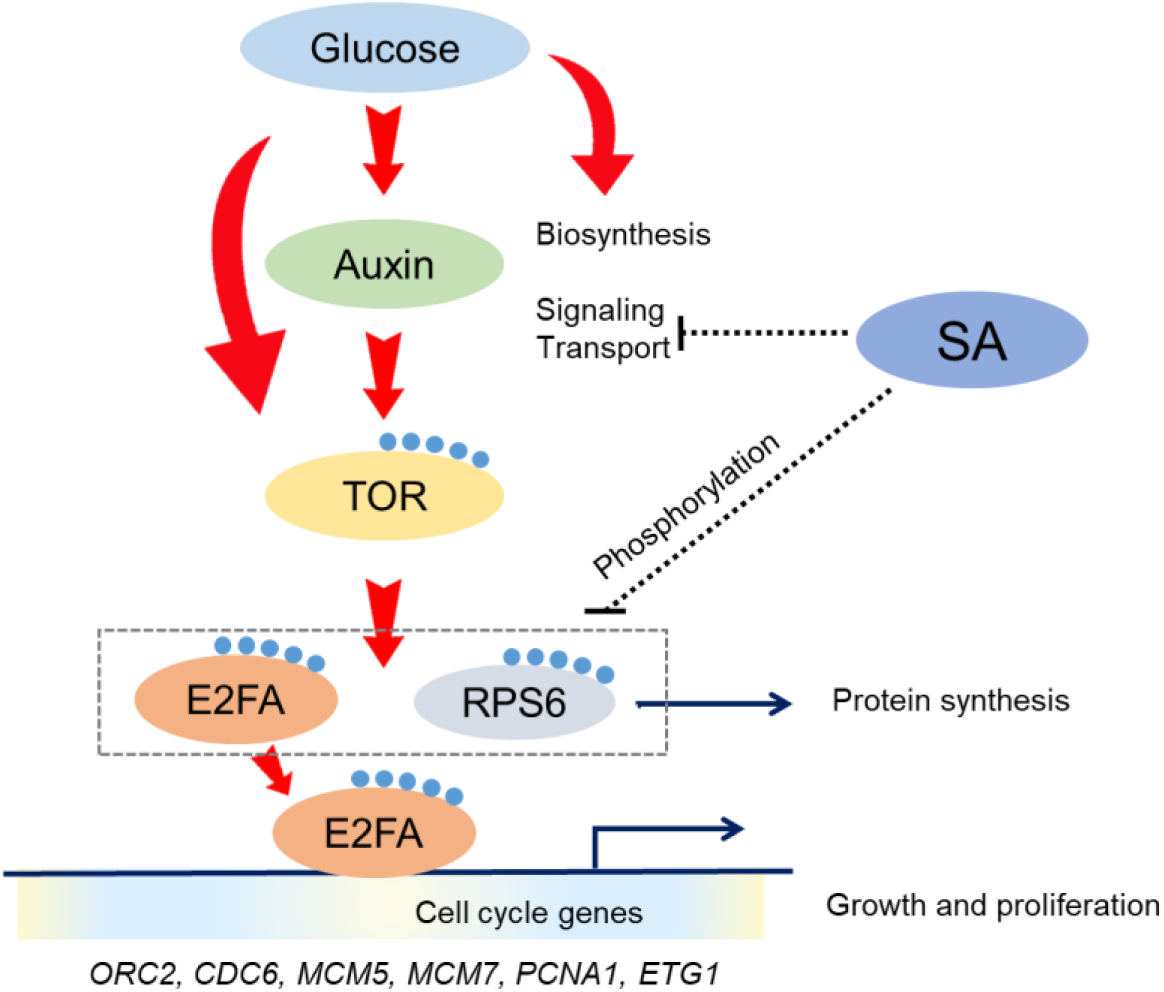
The interplay of glucose, auxin and TOR-E2Fa in mediating root growth repression by SA. A testable model for glucose and SA crosstalk. Glucose promotes root growth by stimulating auxin biosynthesis, signaling and transport. Both glucose and auxin activate TOR kinase, which, in turn, activate RPS6 and E2Fa, key players in cell cycle regulation. E2Fa binds to S-phase cell cycle gene promoters, upregulating their expression and promoting cell growth and proliferation. Conversely, SA appears to inhibit root growth by antagonizing auxin signaling and transport. Additionally, SA reduces the phosphorylation levels of both RPS6 and E2Fa, negatively impacting cell growth and proliferation processes, ultimately leading to root growth inhibition.

## Acknowledgments

SSR acknowledges Council of Scientific and Industrial Research (CSIR) and NIPGR for research fellowships. AL acknowledges NIPGR core grant, DBT: NWBA-2015 (BT/HRD/NWBA/37/01/2015(xii)) and JC Bose fellowship 2021 (JCB/2021/000012). The authors are grateful to the DBT-eLibrary Consortium (DeLCON) for providing access to e-resources. The authors thank Dr. Senthil-Kumar Muthappa for *sweet11/sweet12* mutant line, Dr. Ranjan Swarup for pAUX1::AUX1-YFP, Dr. Christian Meyer for *tor 35-7* mutant line, Dr. Julio Salinas for pNPR1::GUS, Dr. Zhonglin Mou for pNPR1::MYC-NPR1, Dr. Zoltan Magyar for pE2Fa::gE2Fa-3xYFP and Dr. Arp Schnittger for CYCB1;1::CYCB1;1-GFP and Dr. Kalika Prasad for WOX5::GFP lines.

## Author contributions

SSR and AL conceived and conceptualized the study. SSR performed the experiments. AL reviewed and complimented the manuscript.

## Declaration of interests

The authors declare no competing interests.

## Supplementary Data

**Fig. S1** Effect of glucose and SA on columella stem layer formation and WOX5 expression.

**Fig. S2** Quantification of *DII-VENUS* intensity of seeds treated with high SA concentration.

**Fig. S3** The effect of root transport mutant *rcn1* and ethylene signaling components on SA mediated root growth inhibition.

**Fig. S4** PR length measurement of the *npr1,npr3,npr4* mutant after glucose and SA treatment.

**Fig. S5** PR length measurement of the glucose sensor mutant, *hxk1-3* mutant after glucose and SA treatment.

**Fig. S6** EdU staining in roots of the WT and *npr1-2* mutant under glucose treatment. Expression of pCYCB1;1-CYCB1;1-GFP expression upon glucose and SA treatment.

**Table S1** List of primers for qRT-PCR analysis.

## Materials and Methods

### Plant materials and treatments

*Arabidopsis thaliana* seeds were surface sterilized for 10 min in 4% (v/v) sodium hypochlorite, washed with autoclaved distilled water five times, placed at 4 °C for 2 d in the dark and then planted on ½ strength Murashige and Skoog (MS) agar medium (Sigma-Aldrich, St Louis, MO, USA; supplemented with 0.8% (w/v) agar and 1% (w/v) sucrose, pH 5.7) and grown at 22 °C+/- 2°C under 16 h light/8 h dark conditions. For chemical treatments [Salicylic acid, PCIB, NPA, TIBA, IAA, Kynurenine from Sigma], the working solutions of the chemicals were directly added to ½ MS agar medium after the autoclaved agar medium was cooled to below 50 °C and then the medium was poured immediately into Himedia square petri dishes (90mmx 90mm). After sealing with Parafilm (American National Can, Greenwich, CT), the plates with Arabidopsis seedlings were incubated vertically in a controlled environment growth chamber at 22 °C with a 16 h light/8 h dark photoperiod (light intensity = 60 µmol m^−2^ s^−1^).

T-DNA insertion and EMS mutants in the Col-0/ Col-1/ Ler/ Ws-2 background for the following genes were obtained from NASC (Nottingham Arabidopsis Stock Centre, Loughborough, UK) and ABRC (Arabidopsis Biological Resource Centre). *npr1-2* (AT1G64280, CS3801), *npr4-2* (AT4G19660, SALK_098460), *npr1-1; npr3-1; npr4-3* (CS72350), *axr1-3* (AT1G05180, CS3075), *axr2-1* (AT3G23050, CS3077), *eir1-1* (AT5G57090, CS8058), *pin3-4* (AT1G70940, CS9363), *pin4-2* (AT2G01420, CS9368), *pin7-2* (Friml et al, 2003), *mdr1-1* (AT3G28860, Noh et al., 2001), *lax3* (AT1G77690; Swarup et al., 2008), *aux1-7* (AT2G38120, CS3074), *eto1* (AT3G51770, CS3072), *etr1-1* (AT1G66340, CS237), *ein2-1* (AT5G03280, CS3071), *ein3-1* (AT3G20770, CS8052), *rcn1-1* (AT1G25490, CS3875), *hxk1-3* (AT4G29130, CS69135), *snc1* (AT4G16890, CS699080), *cpr5-2* (AT5G64930, CS3770), *PIN2::PIN2-eGFP* (NASC), *tor RNAi 35-7* (Deprost et al., 2007), DII-VENUS (Brunoud et al., 2012), *sweet11/12* (AT3G48740, AT5G23660, CS68845), *NPR1::GUS* (Dr. Julio Salinas), *NPR1::MYC-NPR1* (Dr. Zhonglin Mou), *PIN1::PIN1-GFP* (NASC), *AUX1::AUX1-YFP* (Dr. Ranjan Swarup), *CYCB1;1::CYCB1;1-GFP* (Dr. Arp Schnittger), *WOX5::GFP* (Dr. Kalika Prasad), TAA1p:GFP-TAA1 (CS16432), pE2Fa::gE2Fa-3xYFP (Prof. Zoltan Magyar).

### RNA extraction

Five-day-old Arabidopsis seedlings grown on ½ strength MS agar medium in a vertical position were transferred to liquid ½ MS medium without sugar for 24 hours in dark (starvation) and then on media containing different concentrations of SA with or without glucose as indicated in the main text. Total RNA was extracted from nearly 100 mg roots of Arabidopsis seedlings according to the manufacturer’s protocol (Qiagen). Total RNA was treated with RNase-free DNase I (Qiagen) to digest genomic DNA contamination; the concentration of the DNaseI-treated total RNA was checked using spectrophotometry.

### cDNA synthesis and quantitative real-time-PCR

First-strand cDNA was synthesized with 2 µg of total RNA using Takara (PrimeScript 1st strand cDNA Synthesis Kit). After 20x dilution with water, 1 μL cDNA was used in 10 μL PCR mixture for quantitative real-time (qRT)-PCR. *UBQ10* was chosen as an internal control gene. Each biological sample was performed with at least three technical repetitions and the data analysis shown was carried out using three independent biological replicates or as indicated in figure legends.

### Immunoblotting assays

Plants were harvested, frozen and ground into fine powder with the help of TissueLyser II (Qiagen) in liquid nitrogen in a microfuge tube. The ground powder was thawed in protein extraction buffer containing 25 mM TRIS-HCl pH 7.6, 15 mM MgCl2, 15 mM EGTA, 75 mM NaCl, 60 mM β-glycerophosphate, 1 mM dithiothreitol (DTT), 0.1% NP-40, 0.1 mM Na3VO4, 1 mM NaF and 1 mM phenylmethylsulphonyl fluoride (PMSF) protease inhibitor. Plant crude proteins were then centrifuged at 13000g at 4°C for 10 minutes and this step was repeated until the supernatant was clear of any debris. For denaturing protein samples, 6X SDS-PAGE protein loading buffer was added to each sample and boiled for 10 min. The protein samples were separated by SDS-PAGE and transferred onto the Nitrocellulose membrane (Amersham) for immunoblot analysis. Washing was done with Tris-Buffered Saline (TBS) containing 0.1% Tween-20. Primary antibodies were used in a 1:10000 dilution-Anti-GFP (Abcam, ab290), Anti-RPS6A-P (Agrisera, AS194302), Anti-RPS6A (Agrisera, AS194292), Anti-MYC (CST, 2278T), p-Thr/Phosphothreonine Antibody (Santa Cruz, sc-57562) and p-Ser/Phosphoserine Antibody (Santa Cruz, sc-81514). Immunoblots of targeted proteins were visualized using HRP-conjugated secondary antibodies with 1:10000 dilution after incubation with appropriate primary antibodies.

### Immunoprecipitation

For IP protocol, Arabidopsis seedlings expressing pE2Fa::gE2Fa-3xYFP were harvested and their proteins isolated (as described above). Protein extracts were mixed and incubated for 1h at 4°C with protein A/G agarose beads (G-Biosciences) for pre-clearing. Around 2 ml of protein sample was then incubated with 1µl of Anti-GFP antibody (Abcam 290) overnight (4°C) on a rotating wheel. pE2Fa::gE2Fa-3xYFP protein was captured using 50 µl protein A/G agarose beads (G-Biosciences) and kept for 4 hours on a rotating wheel at 4°C. Beads were collected by centrifugation (3800g, 4°C) and washed with 1XPBS buffer thrice. Supernatant was discarded and 2X laemelli buffer was added to the beads. After brief heat denaturation at 95°C on heat block, the samples were run on SDS PAGE gel. Western blotting was performed as described in previous sections.

### Measurements and statistical analysis

Five-day-old light-grown seedlings were transferred to treatment medium and their root tips were marked at the back of the square petri dish. Digital images were captured after 7 days using a Nikon Coolpix digital camera. Total root length of the seedling after transfer was measured using the ImageJ program from the National Institutes of Health. A minimum of three independent biological experiments were performed until otherwise indicated in figure legends. Values were statistically analyzed by ANOVA or by Student’s t-test wherever indicated. Different alphabets indicate significant differences at *P < 0.05*.

### Laser confocal scanning microscopy

Leica TCS SP8 AOBS Laser Confocal Scanning Microscope (Leica Microsystems) was used throughout the study. Roots were stained in 10 μg/ml propidium iodide (PI) and immediately observed under the eyepiece. The laser, pinhole and gain settings of the confocal microscope were kept identical among all treatments. PI and GFP were detected with a band-pass 570-670 nm filter and 500-545 nm filter, respectively. For the yellow fluorescent proteins (YFP)-tagged reporters, the excitation wavelengths were 488 nm and fluorescence were collected in the ranges of 493-536 nm. All experiments were repeated at least three times or as indicated in figure legends.

### EdU Staining

Seeds of Arabidopsis were sown on ½ MS media without any sugar for four days and covered with a foil to exhaust photosynthetic sugar, after which they were treated with liquid media supplemented with either glucose or no sugar (control). 10 µM EdU (Invitrogen Click-iT® EdU Imaging Kit) was then added to a final concentration and the samples were kept on an incubator shaker maintained at 22°C. The seedlings were then fixed in freshly prepared fixative solution containing 3.7% (v/v) paraformaldehyde (Sigma) and 1% (v/v) Triton-X 100 (Sigma) in 1×PBS solution for 1h and washed twice with 3% (w/v) bovine serum albumin (Himedia) in 1×PBS solution. The seedlings were then incubated with 50 µl Click-iT® reaction cocktail (Invitrogen Click-iT® EdU Imaging Kit; 43 µl of 1 × Click-iT® EdU reaction buffer, 2 µl of CuSO4, 0.12 µl of Alexa Fluor® azide and 5 µl of 1 × Click-iT® EdU buffer additive) for 1 h protected from light at room temperature. The stained seedlings were then washed once with 3% (w/v) BSA in 1 × PBS solution and then stored in 1 × PBS solution protected from light until imaging. Seedlings were mounted in 1×PBS solution and imaged using Leica TCS SP8 confocal microscopy. The excitation wavelength used was 647 nm and emission wavelengths between 655 and 750 nm were collected. Differential interference contrast (DIC) images of the seedlings were also captured in addition to the EdU stain.

### Histological GUS staining

For GUS staining, tissues were immersed in GUS staining solution (50 mM sodium phosphate buffer, pH 7.5, 0.1% (v/v) Triton X-100, 2 mM Potassium ferricyanide, 2 mM Potassium ferrocyanide and 1mg/ml X-Gluc-5-bromo-4-chloro-3-indolyl-b-glucuronic acid) at 37°C and stopped after the signal was achieved. Seedlings were cleared and kept in 50% (v/v) ethanol, 10% (v/v) acetic acid until photographed.

## Notes

### Competing Interest Statement

The authors have declared no competing interest.

